# Correlates of cannabinoid concentrations, real-world driving, and driving-related skills

**DOI:** 10.1101/387936

**Authors:** Mark B. Johnson

## Abstract

Research on the relationship between cannabis use and safe driving has produced mixed results. Most studies have focused exclusively on the presence or concentration of delta-9-tetrahydrocannabinol (THC), the primary psychoactive ingredient in the drug. However, cannabis is a complex substance, and both toxicological research and user experience suggests that some cannabis strains—often those with at least moderate levels of *cannabidiol* (CBD)— produce a different, more sedating “high” than cannabis strains with no or low levels of CBD. We hypothesize that the sedating properties of some high-CBD cannabis strains has potential to impair driving and driving-related skills above and beyond the effects of THC intoxication. Three studies—one instrumented vehicle driving study and two laboratory-style epidemiological studies—examined real driving and computerized task performance as a function THC and CBD concentrations (and their interactions). In all three studies, higher CBD levels predicted greater impairment. There was relatively little evidence of impairment when CBD was zero, even at high THC levels. The results suggest that THC concentrations alone are not sufficient to predict impairment due to cannabis use. Results are interpreted in the context of drug tolerance.

## Introduction

### Current state of the research

Recent legalization of cannabis has spurred interest in the relationship between acute cannabis use and driving impairment. In fact, drug-involved driving appears to be on the rise [1-4]. Despite this increased focus, it remains unclear whether and to what extent cannabis contributes to crash involvement.

On one hand, experimental cannabis dosing studies routinely find evidence of dose-specific impairment on cognitive and psychomotor driving-related skills, as well as simulated driving [5, 6]. On the other hand, epidemiological studies that predict actual vehicle crashes from delta-9-tetrahydrocannabinol (THC) measured in drivers’ systems have been inconsistent [7, 8]. A recent meta-analysis [9] reported that approximately one in three studies examined found no association between driving under the influence of cannabis and crashes.

### Delta-9-tetrahydrocannabinol (THC) and cannabidiol (CBD)

One limitation of the extant research on cannabis and driving impairment is that the field has failed to consider the complexity of the drug. Cannabis contains over 113 identified cannabinoids [10] and many more terpenes and flavonoids; yet researchers have focused exclusively on THC as the sole predictor of user experience and the agent of driving impairment. There is growing understanding that distinct subspecies of the plant, *cannabis sativa* and *cannabis indica*, can produce qualitatively different types of “high.” Sativa strains are associated with producing a more energetic, cerebral high, while indica strains often are associated with more sedative effects [11-13].

It may be possible to predict whether cannabis is excitatory or sedating by the composition of its constituent compounds THC and CBD. Energizing sativa strains consumed by users often are characterized by higher concentrations of THC and very low levels of the CBD, whereas sedating indica strains often contain a more balanced mixed of each [14]. Whereas THC is responsible for the euphoric high of cannabis [15], some studies suggest that CBD calms the negative THC side effects of anxiety and paranoia [16, 17], while other studies directly link cannabis high in CBD to mental and physical sedation [12, 18, 19]. This pharmacological research on high-CBD cannabis dovetails somewhat with the experiences reported by cannabis connoisseurs who describe cannabis high in CBD mixed with THC as producing a sleepy, dreamlike experience and cannabis high in CBD alone as producing a lethargic “body-stoned” experience [20, 21].

The relationship among cannabis strains, THC/CBD concentrations, and behavioral effects is notably complex. For example, Russo [22, 23] argues that the sedating effect associated with indica cannabis is not due to CBD, but rather due to the terpene *myrcene*, which happens also to be prevalent in indica. Accordingly, the relationship between high CBD cannabis and sedation is argued to be spurious. However, even as a proxy variable, CBD may be useful in distinguishing cannabis that produces sedating versus euphoric experiences. This research does not attempt to make causal pharmacological claims—it only attempts to improve prediction by measuring and modeling CBD in addition to THC.

The fact that some cannabis strains induce lethargy and sedation while other strains have more heady and excitatory effects may have implications for driving, which heretofore have not been explored. Given that attention and vigilance are key factors to safe driving [24, 25], anything that promotes sedation might interfere with one’s ability to maintain focus on the roadway, even if it does not make one intoxicated or high. Following the assumption that CBD might be a useful epidemiological measure to distinguish sativa versus indica stains, we wish to examine whether CBD can be used to predict driving impairment independently from THC. Described herein are three studies that provide preliminary evidence that THC alone is insufficient to predict driving impairment, and an investigation of compounds other than THC is needed to better explain the public health risks of cannabis use.

### Obstacles to cannabis research

Conducting research on the effects of cannabis consumption of human behavior can be challenging. In the United States, cannabis remains on the Schedule I list of drugs, greatly limiting who can obtain, manage, and administer the drug for research purposes. Furthermore, even for those qualified, the cannabis legally available is limited to that grown under the Drug Supply Program of the National Institute on Drug Abuse (NIDA). While the menu of NIDA’s Drug Supply Program has expanded somewhat in recent years, strains that are very high in THC, CBD, or both were not available at the time this research was conducted.

For these reasons, the research described herein does not include a formal controlled dosing study, as such could not be legally conducted using the strains required to address the research questions. Rather the studies are epidemiological in nature, where objective information on cannabis consumption (quantitative concentrations of THC and CBD) is obtained through biological assays of participants’ oral fluid. One study is a naturalistic driving study where participants’ THC and CBD concentrations are used to predict vehicle performance; the two other studies are laboratory-style studies of driving-related skills where participants self-dose with varied strains of cannabis. Strengths and limitations of these studies are discussed.

## Study 1

### Study 1: Materials and methods

The Institutional Review Board of the Pacific Institute for Research and Evaluation approved all aspects of the Study 1 research described herein.

#### Recruitment

Participants were recruited via advertisements in cannabis dispensaries and through listservs of private cannabis clubs in the Denver, Colorado, area. Interested individuals were directed to an online prescreening survey, which included items on cannabis and alcohol use, driving practices, and demographics. Also included was the Drug Abuse Screening Test (DAST) [26] and a brief 7-item psychosis screener [27].

Eligible participants (a) used cannabis at least twice monthly; (b) drove several times per week; (c) were aged 21 and older; (d) were not pregnant; (e) had no more than two moving violations or one at-fault accident in the past 3 years; (f) had no driving while intoxicated (DWI) or driving under the influence (DUI) arrests on their driving record; (g) received a score of 12 or lower on the DAST; (h) had no use of illicit drugs other than cannabis; and (i) had no indication of psychosis on the psychosis screener. All eligible individuals were contacted by phone by the project manager for further assessment. To be invited to participate, individuals also needed to have regular access to a vehicle and serve as the sole driver of that vehicle for up to 10 days during data collection. Those who still were interested were scheduled for an informational and training session.

#### Protocol

The study involved two basic tasks: (1) allowing one’s vehicle to be equipped with instrumentation and driving normally for up to 10 days; and (2) self-collecting oral fluid and breath samples during each driving trip.

##### Instrumentation

Participants’ vehicles were equipped with Aaronia GPS data loggers (http://www.aaronia.com/products/spectrum-analyzers/GPS-Logger). The devices recorded vehicle GPS information (e.g., coordinates, speed, heading, etc.) at one reading per second. The devices also recorded acceleration data at four readings per second. These data indicated change along the X, Y, and Z-axes, where more severe and abrupt car movements translated into larger g-force measurements. Along the X-axis, positive scores reflected faster acceleration, while negative scores indicated harder braking. Along the Y-axis, larger scores (in absolute value) reflect harder turning and swerving to the right and left. Z-axis scores (rising and falling) were not analyzed.

##### Biological samples

Participants were instructed to provide one or more oral fluid samples, using Quantisal^™^ collection tubes, each time they went driving. Because THC concentrations in oral fluid are highly volatile during the first hour, participants were asked to provide a second sample after 30 minutes of driving if they had used cannabis within 60 minutes before the start of their trip. Participants were instructed to label each oral fluid sample with the date and time.

Oral fluid samples were assayed for THC and CBD by Immunalysis Corporation (Pomona, California). There was no option to directly assess myrcene levels from oral fluid. Confirmation tests (to obtain quantitative concentrations) were performed using gas chromatography-mass spectrometry (GC/MS) or liquid chromatography-mass spectrometry (LC/MS/MS) technology. This method for assaying oral fluid for THC has a limitation of quantification of 1 ng/ml, linearity of 0.5 to 32 ng/ml, intraday precision of 7.1% at 3 ng/ml and 2.9% at 12 ng/ml, and interday precision of 4.9% at 3 ng/ml and 1.6% at 12 ng/ml. The same laboratory and methodology were used to assay samples for the National Highway Traffic Safety Administration’s National Roadside Surveys [28, 29]. In addition to oral fluid collection, participants were given a calibrated breathalyzer that did not display results (but rather stored results internally) and were instructed to provide a breath sample each time they provided a saliva sample.

##### Driving

After participants were trained and instrumentation was installed, participants were asked to drive normally for 6 to 10 days. Research assistants met with each participant every other day to collect their oral fluid samples.

#### Data elements

##### GPS data

Time-stamp GPS readings provided a temporal framework for the dataset. A “driving trip” was defined as 10-minute break with the vehicle not moving. Over the course of the research, participants took between 2 and 43 driving trips (median = 17). GPS coordinate data were geocoded to reflect individual roads and formal road classifications (i.e., parking lot, interstate, expressway, arterial roadway, residential street, and ramps/exist). The GPS sensor provided information on vehicle speed and heading.

##### Accelerometer data

Accelerometer data were collected at 4 readings per second and were automatically linked with GPS data (see above). The study’s main dependent measures were created from X- and Y-axis acceleration scores. However, these went through several stages of processing before being analyzed. First, acceleration scores were centered on the mean score of each driving trip. Some participants removed the instrumentation from their vehicles between driving trips and reinstalled them at different pitches and yaws, which appeared to have introduced trip-level bias into scores. Mean centering for each trip accounted for that.

Second, the acceleration data was aggregated into 5-second blocks (see below), which served as the unit of analysis for the study. From the 20 scores within each 5-second block (5 seconds x 4 readings per second) we computed the dependent measures.

##### Dependent measures

The study focused on three dependent measures—elevated g-force events, standard deviation of acceleration, and vehicle speed. Except for vehicle speed, separate values were computed for X- and Y-axes. First, a dichotomous *elevated g-force event* value (0 or 1) was computed for each 5-second block of driving data (see Final Data Structure, below). Prior research found that “jerky driving”—for example, driving events (such as braking) exceeding 4.0 m/s^2^ (.408 gs)—predicted crashes and near crashes [30, 31]. Accordingly, 5-second blocks that included absolute accelerometer readings of .408 gs or greater were coded as “elevated.”

Second, we computed the *standard deviation* of raw acceleration scores with each 5-second block (after first removing outliers ± 3 standard deviations [SDs]). This outcome was interpreted as reflecting lack of longitudinal (X-axis) and lateral (Y-axis) control—that is, less stable speed maintenance and lane position. Third, we computed the mean *speed* for each 5-second block.

##### Cannabinoid and alcohol concentrations

Assays of oral fluid produced quantitative concentrations (in ng/ml) of THC and CBD. We determined *a priori* that drug concentrations would be valid for a 10-minute period and used to predict driving behavior within that period. If an oral fluid sample was collected at 1:05 p.m. on September 5^th^, those resultant drug concentrations would be used to predict only the driving behaviors that occurred from 1:00 to 1:10 p.m. on that day. The decision to limit that analysis to a 10-minute window was made to maximize sensitivity while retaining adequate sample size. There was almost no cooccurrence of mixing alcohol with cannabis. There were very few positive breath sample readings in this sample, and none greater than .02 g/dl. Therefore, alcohol readings were not included in analysis.

#### Final data structure

The final data structure is depicted in Fig 1. Participants took multiple driving trips. Trips included one or more 10-minute blocks for which quantitative THC and CBD concentrations were available. Within each 10-minute block there were 120 5-second blocks of accelerometer driving data (these 5-second blocks were the unit of analysis). We did not examine driving data that occurred outside the 10-minute blocks for which we had cannabinoid data. Dependent measures reflected driving behavior during these 5-second intervals, including elevated X- and Y-axes g-force events, X- and Y-axis standard deviations, and mean speed. These driving outcomes were predicted to vary as a function of THC and CBD concentrations. This focus on small-units of driving behavior is a departure from other instrumented-vehicle studies [30] that used whole trips as the unit of analysis.

**Figure 1.**
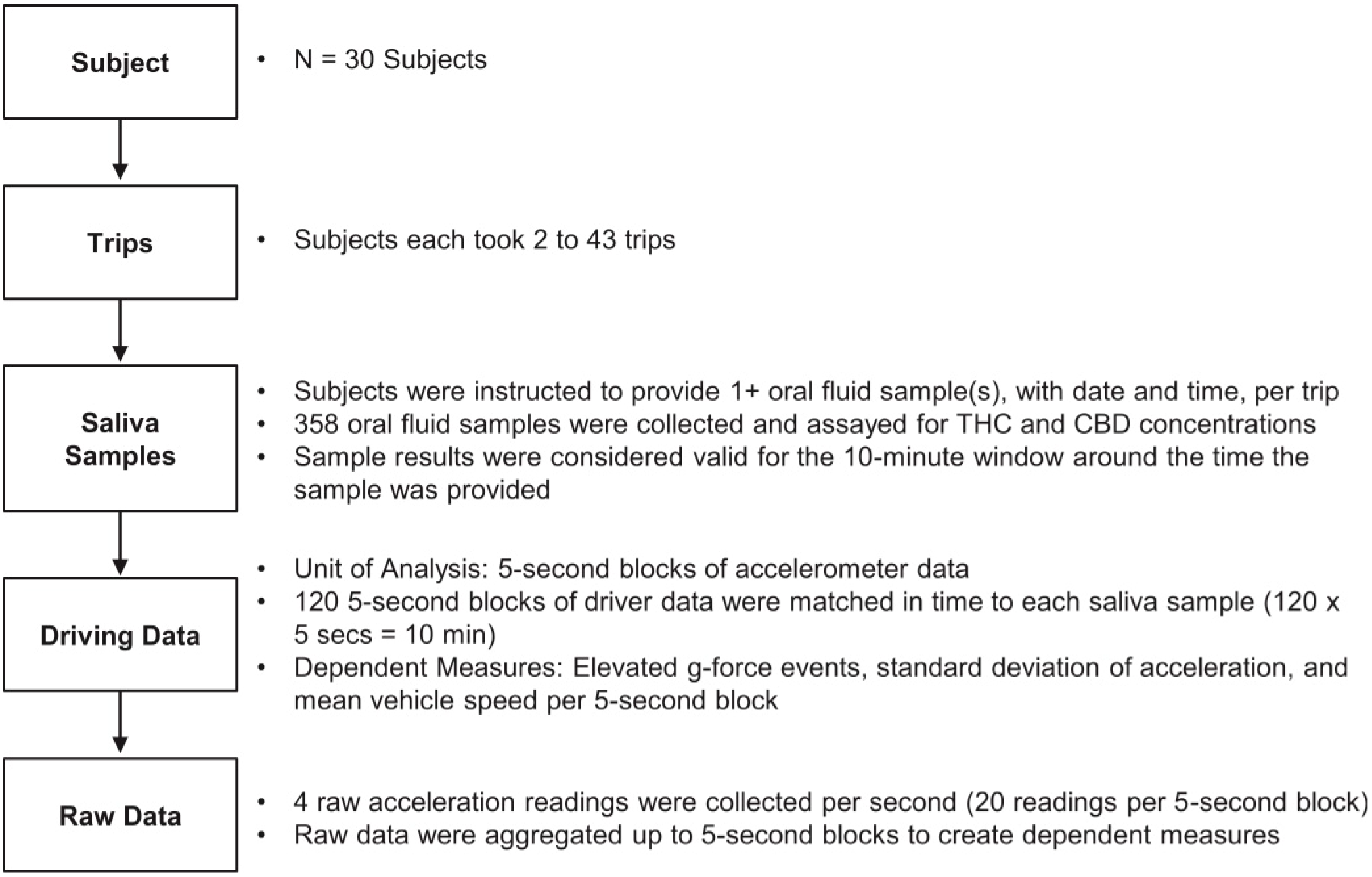
Final Structure of IVDIRM Data

The analytic goal was to use THC and CBD concentrations to predict driving behavior. We used generalized linear mixed modeling (GLMM) in SAS 9.2 to predict driving behavior from THC and CBD. We accommodated the multiply-nested data structure, modeling subject (R-side) and trip-within-subject (G-side). This accounted for that fact that each driver produced many data points, and accelerometer scores may have been influenced not only by cannabinoids and individual driver characteristics, but also by features of the specific driving trip and types of roads being driven. Driver characteristics and road type were controlled for as well as additional fixed effects.

## Study 1: Results

### Sample

A total of 91 individuals completed the online prescreening survey; 40 were deemed eligible and 30 consented to take part in the driving study. These 30 regular cannabis users allowed the installation of instrumentation in their vehicles and provided self-administered oral fluid samples during driving trips over a period of 6 to 10 days. A slight majority (56.7%) was male. Ages ranged from 22 to 57 (median = 37). Most of the sample was White, non-Hispanic (63.3%), with five Black, four Hispanic, one Asian, and one Native American driver.

Most participants (60%) used cannabis several times per day, while the remainder used daily or almost daily. All participants had a history of driving while under the influence of cannabis. Most participants (70%) drove within 2 hours of using cannabis at least weekly while the remainder did so at least monthly. The majority of participants had not driven within 2 hours of drinking (86.7%) or driving after drinking “too much” (96.7%) in the past six-months; those who had driven after drinking did so only once or twice.

### Data summary

Our sample of 30 drivers took a total of 439 driving trips (median = 17 per driver)— approximately 185 hours of driving. A total of 2.66 million independent acceleration records were captured along the X- and Y-axes. The distribution of dependent measure scores—the proportion of X- and Y-axis elevated g-force events, mean standard deviations of X- and Y-acceleration, and mean vehicle speed—is provided in Table 1. For standard deviation of acceleration scores, outliers (scores that exceeded ± 3 SDs) were removed.

**Table 1.** Descriptions of Dependent Measures Used in Analysis.

| **Dependent Measure** | **Value** | **Standard Deviation** |
| --- | --- | --- |
| Elevated X-axis acceleration events | 0.120 | .002 |
| Elevated Y-axis acceleration events | 0.063 | .002 |
| Standard deviation of X-axis acceleration | 0.080 | .040 |
| Standard deviation of Y-axis acceleration | 0.080 | .031 |
| Mean vehicle speed | 27.2 | 16.5 |

Participants produced 358 valid and matched saliva samples from 258 distinct driving trips (1.4 samples per trip); thus, not all driving trips had a matching saliva sample. Only a small proportion of oral fluid samples (10.9%) tested negative for THC. The median THC concentration was 157 ng/ml, while the mean was 454 ng/ml (SD = 721.3). Most CBD concentrations (67.3%) were 0 ng/ml. The mean CBD value was 3.0 ng/ml (SD = 20.1) but the maximum was 290 ng/ml. THC and CBD concentrations were significantly, but not strongly, correlated, *r*(358) = .27, *p* < .01.

Given that the distribution of THC scores was positively skewed, we subjected those values to a natural logarithm transformation (adding .0001 to all cases to make it possible to solve the 10.9% of cases where THC was 0). Because two-thirds of the oral fluid tests were negative for CBD and the positive scores were highly skewed, it was unclear whether any quantitative transformation was appropriate. Therefore, we decided to dichotomize CBD (CBD = 0 ng/ml or CBD > 0 ng/ml) for the analyses.

### Predicting driving behavior for cannabinoid concentrations

Analysis focused on elevated X- and Y-axis events, the standard deviation of X- and Y-axis acceleration, and vehicle speed. The primary predictors were the natural log (ln) THC concentrations, CBD category (0 versus >0), and the ln THC x CBD category interaction. Driver sex, race (White versus non-White), age, frequency of cannabis use, and road type were covariates. Subject and trips-within-subject were modeled as random effects.

<u>Elevated X-axis events</u> involved observed occurrences of driving where acceleration or braking exceed .408 g-forces (4.0 m/s^2^). Analysis used generalized linear mixed modeling to predict the likelihood of elevated X-axis events from cannabinoid concentrations, controlling for driver characteristics, cannabis use frequency, road type, and random effects. Results of analysis are displayed in Table 2.

**Table 2.**
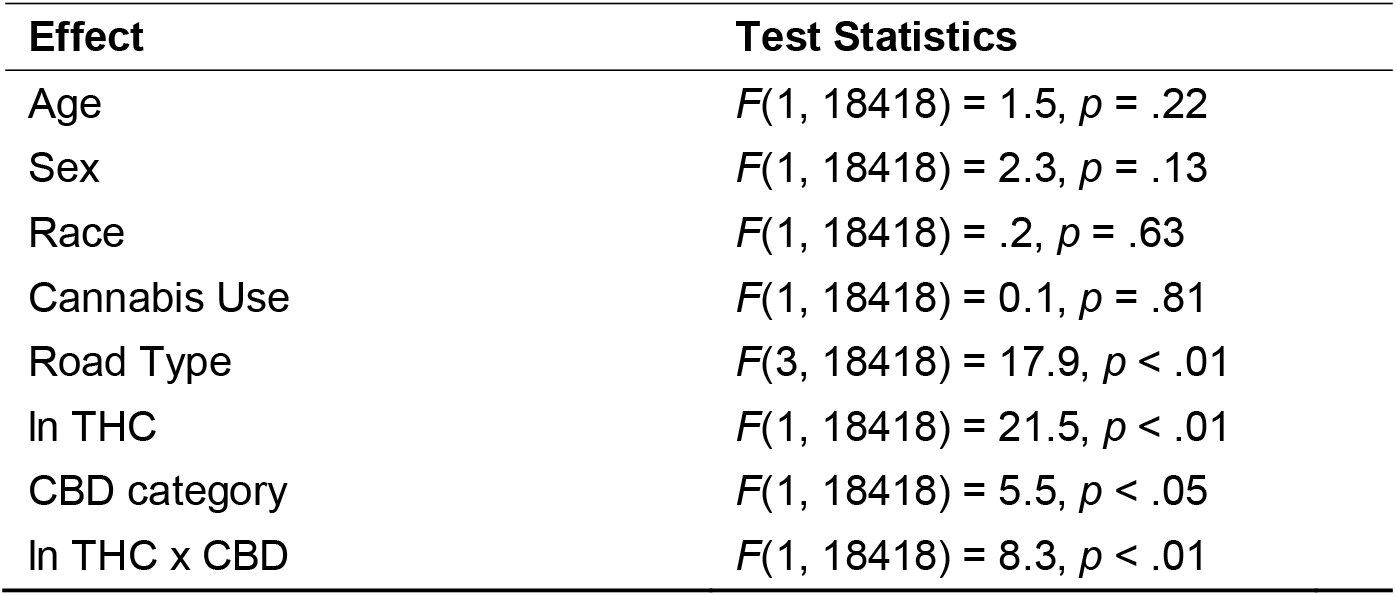
Analysis of the Likelihood of Elevated X-Axis Events.

| Effect | Test Statistics |
| --- | --- |
| Age | $F(1, 18418) = 1.5, p = .22$ |
| Sex | $F(1, 18418) = 2.3, p = .13$ |
| Race | $F(1, 18418) = .2, p = .63$ |
| Cannabis Use | $F(1, 18418) = 0.1, p = .81$ |
| Road Type | $F(3, 18418) = 17.9, p < .01$ |
| ln THC | $F(1, 18418) = 21.5, p < .01$ |
| CBD category | $F(1, 18418) = 5.5, p < .05$ |
| ln THC x CBD | $F(1, 18418) = 8.3, p < .01$ |

Model-estimated likelihoods were computed across the range of THC scores, separately for samples with 0 CBD and those with CBD > 0. The results pattern is reflected in Fig 2. Here likelihood of elevated acceleration/braking events are linked to increasing THC, with a significantly steeper slope associated with the combination of THC and CBD.

**Figure 2.**
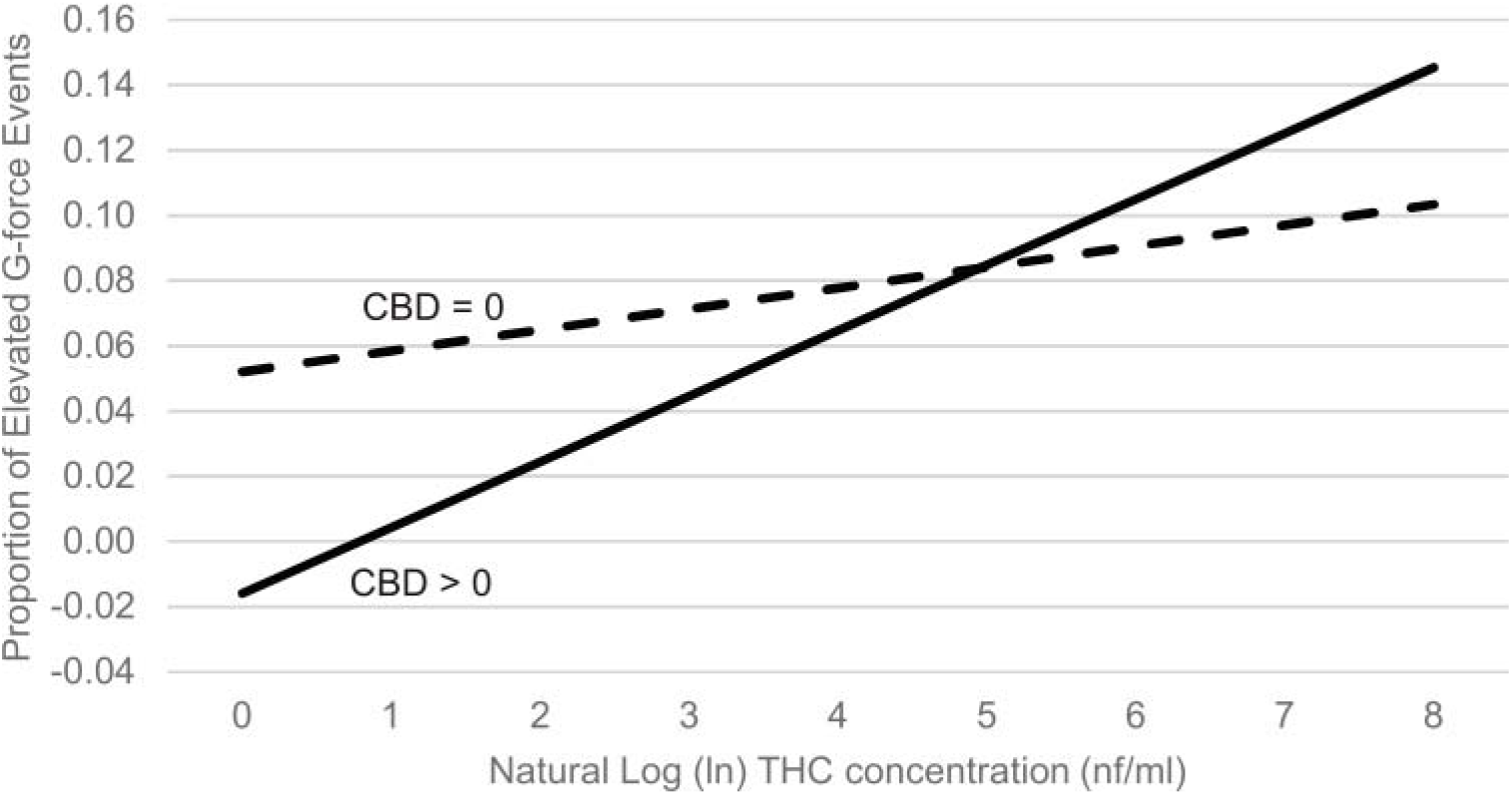
Proportion of Elevated X-Axis G-Force Events as a Function of ln THC and CBD Category

<u>Elevated Y-axis events</u>. Analysis of cannabinoid concentrations on elevated Y-axis events yielded a marginally significant ln THC x CBD interaction (*p* = .06), and we present the pattern of results for descriptive purposes in Fig 3. A main-effects analysis (removing the interaction effect) revealed a significant main effect of CBD category but not ln THC (see Table 3). Accordingly, a significantly lower rate of elevated Y-axis events was observed when CBD category was negative versus when CBD was positive.

**Figure 3.**
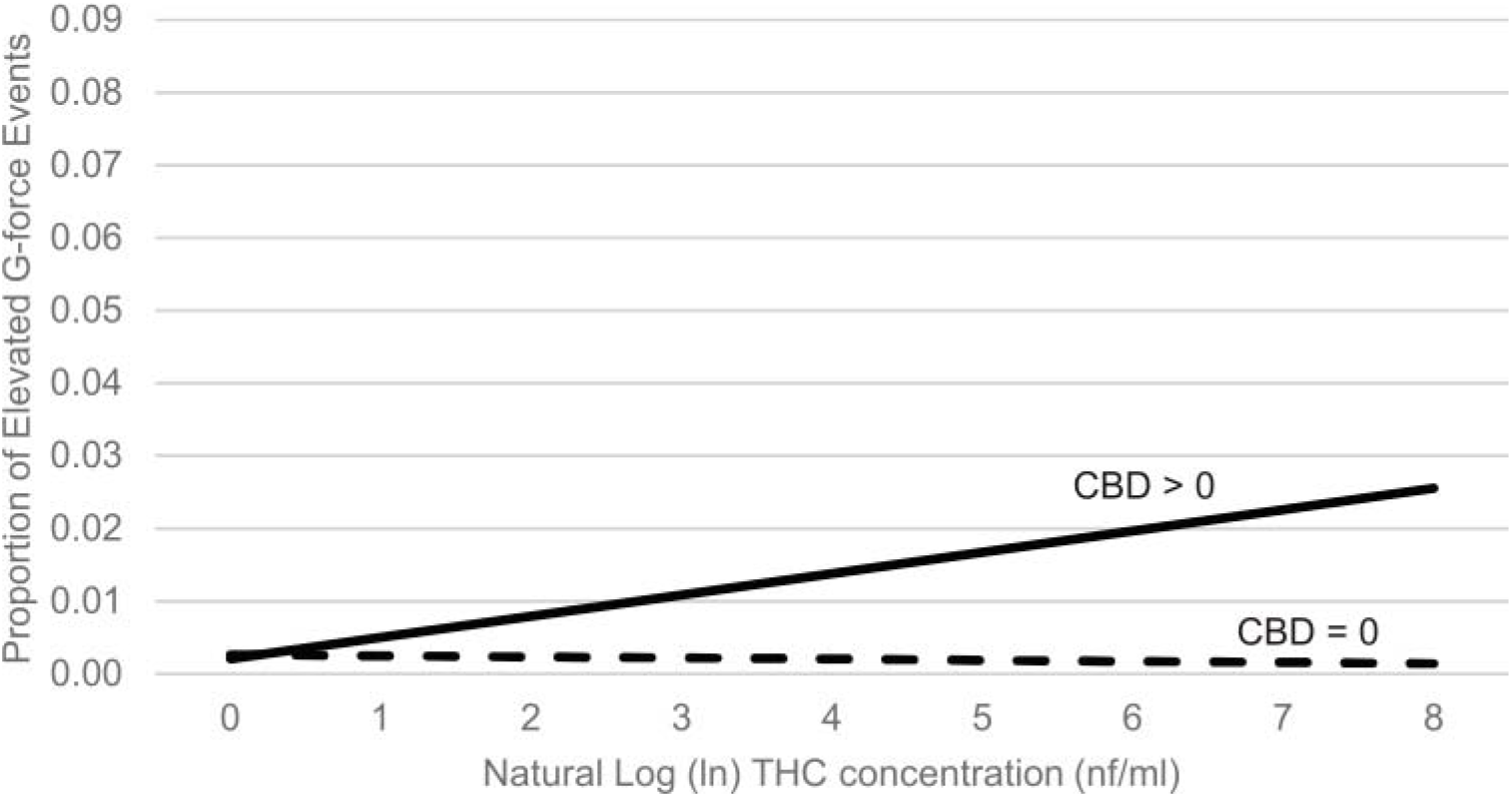
Proportion of Elevated Y-Axis G-Force Events as a Function of ln THC and CBD Category

**Table 3.**
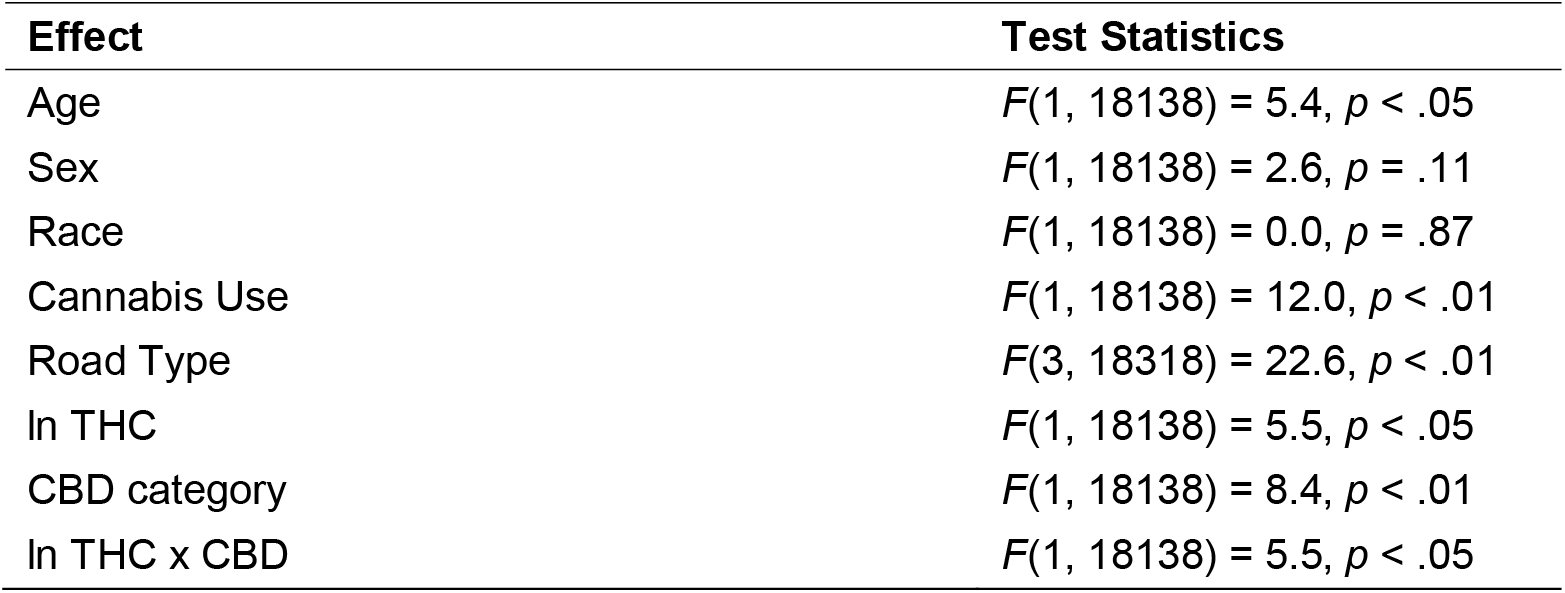
Analysis of the Likelihood of Elevated Y-Axis Events.

<u>X-axis standard deviation</u> acceleration scores reflect the extent to which drivers exhibited less consistency in speed within each 5-second unit of analysis. Outliers were removed. Analysis revealed statistically significant main effects of ln THC, CBD category, and the THC x CBD interaction (see Table 4). Driver age and frequency of using cannabis also were statistically significant.

**Table 4.**
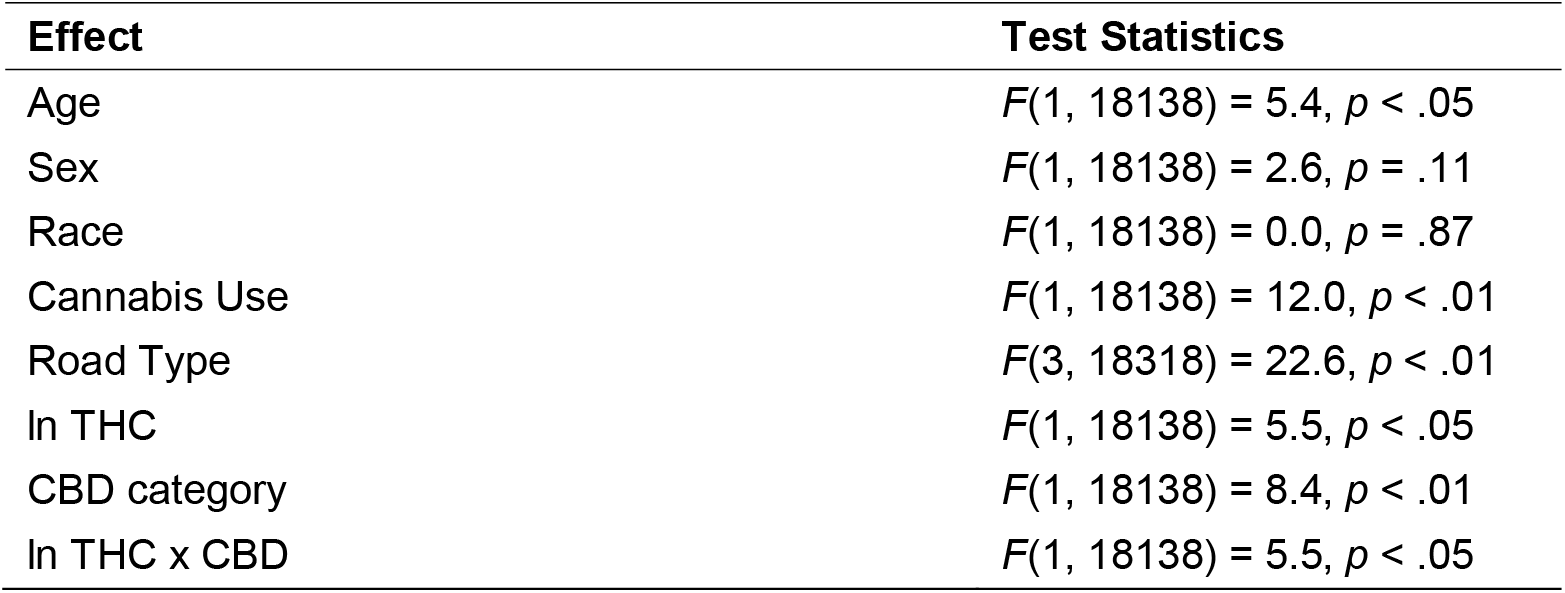
Analysis of the Standard Deviation of X-Axis Acceleration.

The ln THC x CBD category interaction is provided in Fig 4. The pattern is quite different than that observed for elevated g-force events. Whereas previously drivers who were high in ln THC and CBD positive exhibited greater impairment, here drivers who were CBD positive but low in THC demonstrated less control regarding vehicle speed. Drivers who were CBD positive but with relatively low THC concentrations showed more variability in X-axis acceleration, but this difference vanishes as THC values increase.

**Figure 4.**
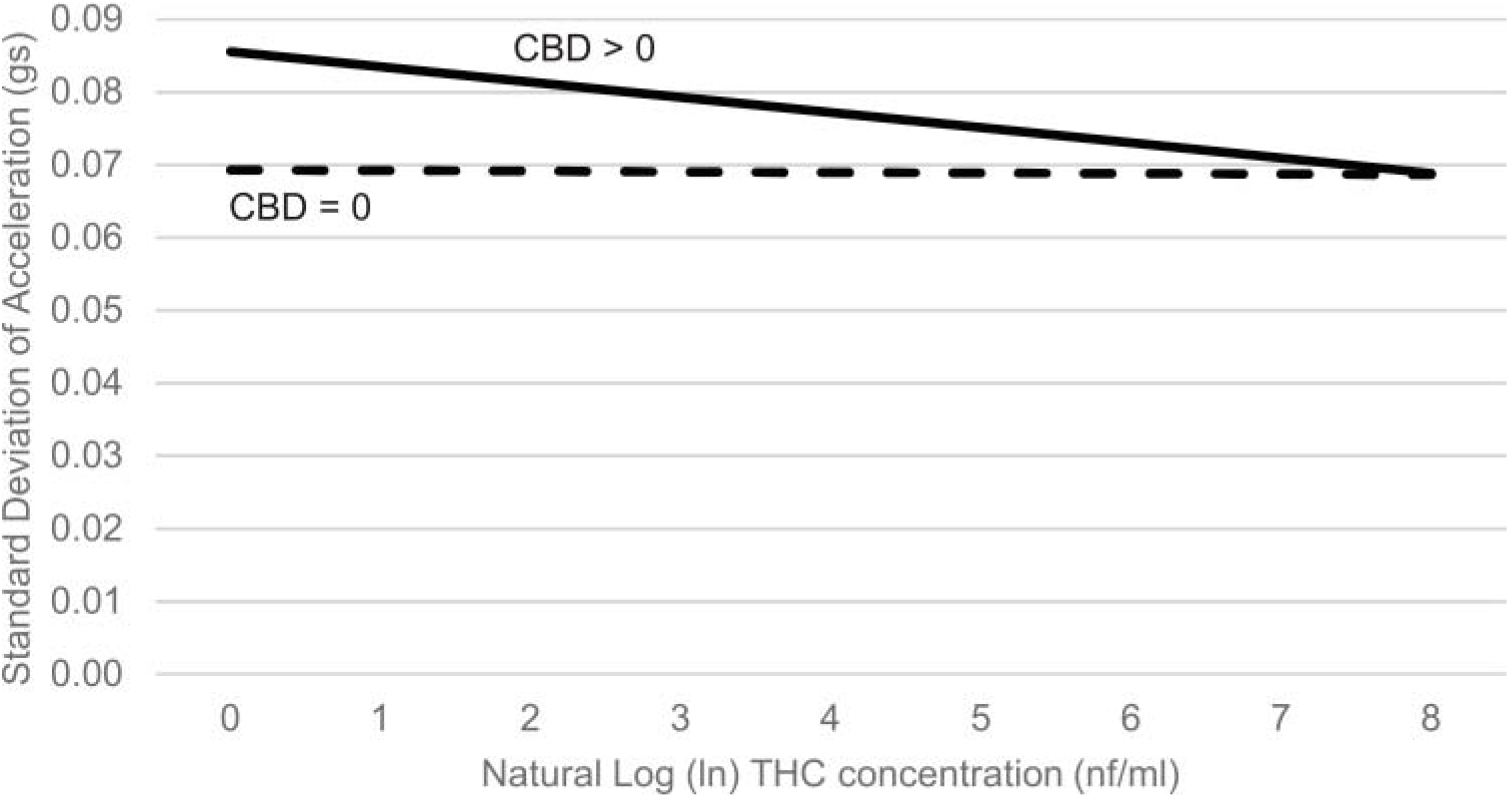
Standard Deviation of X-Axis Acceleration as a Function of ln THC and CBD Category

Older drivers exhibited less variability in X-axis acceleration while driving (B = -0.0007), while drivers who used cannabis relatively more frequently demonstrated greater variability in X-axis acceleration (B = 90.1).

<u>Y-axis standard deviation</u> acceleration scores reflect the extent to which drivers exhibited greater variability in lateral movement (e.g., lane position) within each 5-second unit of analysis. Outliers were removed. Analysis revealed no statistically significant ln THC x CBD interaction (p = .25); however, a main effects model revealed a significant main effect of CBD category (see Table 5).

**Table 5.** Analysis of the Standard Deviation of Y-Axis Acceleration.

| **Effect** | **Test Statistics** |
| --- | --- |
| Age | $F(1, 18188) = 4.0, p < .05$ |
| Sex | $F(1, 18188) = 1.3, p = .35$ |
| Race | $F(1, 18188) = 0.1, p = .81$ |
| Cannabis Use | $F(1, 18188) = 1.4, p = .25$ |
| Road Type | $F(3, 18188) = 40.1, p < .01$ |
| $\ln$ THC | $F(1, 18188) = 3.4, p = .07$ |
| CBD category | $F(1, 18188) = 4.5, p < .05$ |

Mean Y-axis acceleration standard deviations were significantly higher in the CBD > 0 category than in the CBD = 0 category (means = .077 gs versus .079 gs), although the difference was quite small in absolute terms.

The main effect of age revealed that older drivers exhibited relatively smaller Y-axis acceleration standard deviations (B = -0.0003).

<u>Vehicle speed</u> was measured via the GPS sensor. There were no speed outliers. Analysis revealed a statistically significant main effect of sex, ln THC and CBD category, as well as the ln THC x CBD interaction (see Table 6). Regression slopes are depicted in Fig 5. Participants with low THC and high CBD drove the fastest, while those with high THC and high CBD drove the slowest. The main effect of sex revealed that men drove slightly faster, on average, than women (means = 32.1 versus 29.9 mph).

**Table 6.** Analysis of the Vehicle Speed.

| **Effect** | **Test Statistics** |
| --- | --- |
| Age | $F(1, 18426) = 2.6, p = .11$ |
| Sex | $F(1, 18426) = 4.9, p < .05$ |
| Race | $F(1, 18426) = 2.4, p = .12$ |
| Cannabis Use | $F(1, 18426) = 0.1, p = .75$ |
| Road Type | $F(1, 18426) = 3187.4, p < .01$ |
| ln THC | $F(1, 18426) = 33.2, p < .01$ |
| CBD category | $F(1, 18426) = 34.4, p < .01$ |
| ln THC x CBD | $F(1, 18426) = 55.5, p < .01$ |

**Figure 5.**
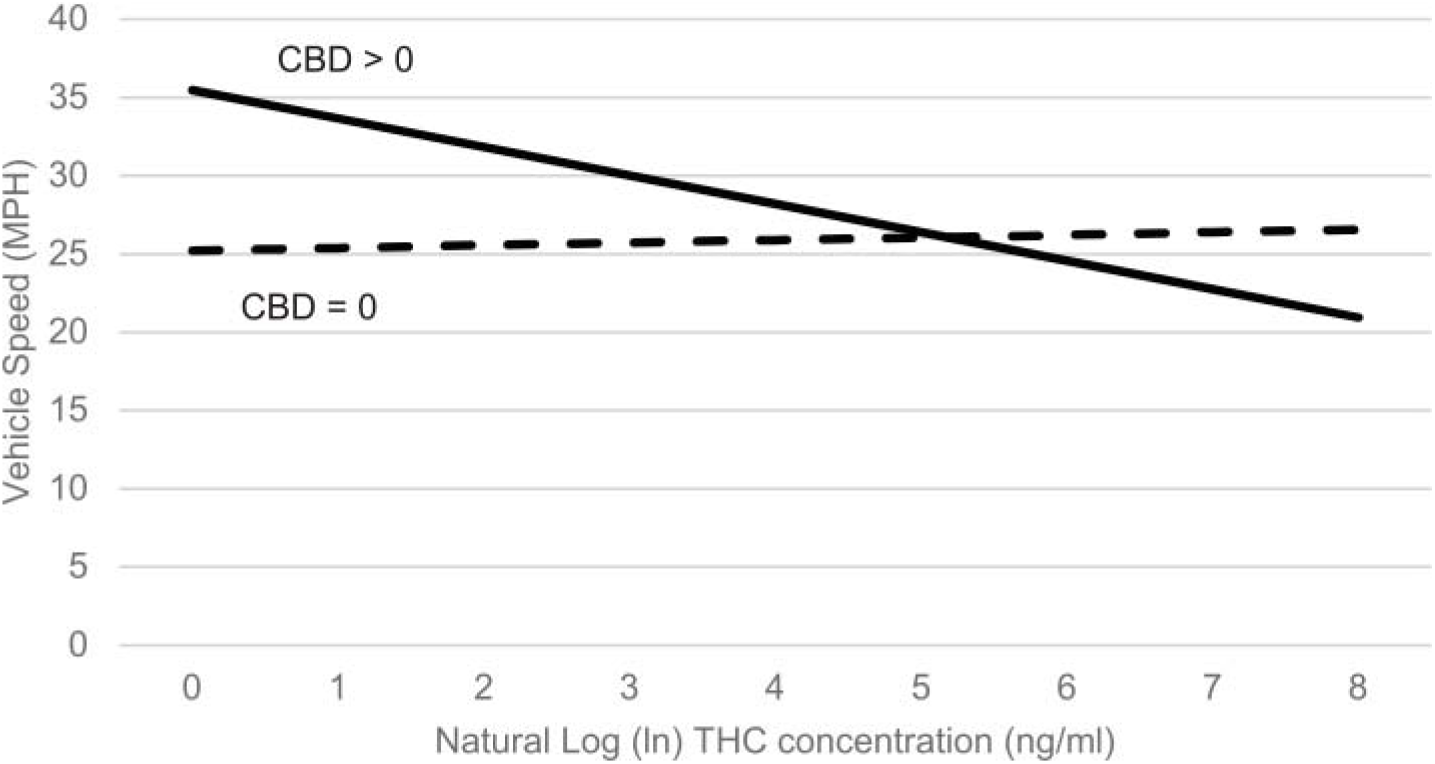
Vehicle speed as a function of THC and CBD category

## Study 1: Discussion

Heavy cannabis users drove instrumented vehicles for several days and provided oral fluid samples that were assayed for concentrations of THC and CBD; drug test results were linked to vehicle performance that occurred within the same 10-minute block as the saliva sample was collected. Because the research examined the same participants over multiple trips and different cannabinoids concentrations, individual personality, driving styles, etc., are accounted for. Evidence that cannabis use predicted driving behavior controlling for individual driver effect is inconsistent with the argument that prior observed relationships between cannabis and risk of crash involvement could be explained by individual differences in risk-taking personality [32].

Cannabis concentrations were statistically related to driving behavior. However, several distinct patterns of results emerged. For elevated g-force events, the combination of being CBD positive and having a high THC was associated with the greatest risky driving for both longitudinal (X-axis) and lateral (Y-axis) movements (although for the latter, the ln THC x CBD category was only marginal, *p* = .06). The relationship between elevated g-force events and THC was weaker when CBD was 0. For the standard deviation of X-axis acceleration and vehicle speed, greatest impairment was observed for drivers who were CBD positive but low in THC. For the standard deviation of Y-axis acceleration, greater impairment was observed for those who tested positive for CBD and was not necessarily related to THC.

In all cases, among the sample of heavy cannabis users, the presence of CBD was important in predicting driving impairment. No other research—experimental or epidemiological—has examined CBD in cannabis as a possible agent of driving impairment. There is research suggesting that CBD attenuates some of the side effects of THC, mainly anxiety and paranoia [16, 17]; it also appears to qualitatively change the drug experience. No results regarding CBD concentrations and driving are in the published literature, but the findings are consistent with evidence that some strains of cannabis are associated with sedation and lethargy and that driving while sedated is a risk factor.

THC by itself (without CBD) was not a strong predictor of impairment. However, previous experimental research found that heavy users do not necessarily show impairment on tests of driving-related skills [33, 34], suggesting that drug tolerance may minimize impairing effects. The participants in this research may have been sufficiently tolerant not to exhibit changes in driving behavior as a function of THC.

## Study 2

The driving study (Study 1) demonstrated, with high ecological validity, that THC alone does not explain the relationship between cannabis consumption and driving impairment; whether participants also tested positive for CBD mattered in a meaningful way. Study 2 was designed as a cost-effective method of further examining the role of THC, CBD, and their interaction, as predictors of impairment using computerized tests of driving-related skills. Study 2 mirrored Study 1 in many ways. Because we could not legally control the strains with which to dose participants, Study 2 relied on participants purchasing their own product, and self-dosing during the laboratory sessions. Lacking *a priori* knowledge and control over the strains participants used, the analyses used concentrations of THC and CBD, assayed from oral fluid, to predict performance on computerized tasks. The design was correlational, therefore, and not experimental. However, note that recently, Hartman and colleagues [35] used a similar correlational approach to relate THC concentrations to performance on a driving simulator (rather than conduct an experimental comparison among low versus high THC strains).

Like all regression/correlational designs, stable estimates require that the methodology produce variability in predictor variable scores as well as on dependent measures. We generated variability from a small sample by using a within-subject design, measuring cannabis concentrations and task performance pre-dosing (baseline) as well as over three post-dosing blocks. Changes in THC and CBD concentrations as the drug is absorbed into the system and then metabolized naturally generated variability, as did differences in potency of the strains used.

### Study 2: Materials and methodology

The Institutional Review Board of the Pacific Institute for Research and Evaluation approved all aspects of the Study 2 research described herein, including the protocol for participants self-dosing with cannabis.

#### Recruitment

Data collection took place over a 4-day period in Denver, Colorado, taking advantage of the legalization of cannabis in Colorado for purchase and recreational use as well as Denver’s robust boutique cannabis market. Participants were recruited via advertisements in cannabis dispensaries and listservs of private cannabis clubs and were directed to an online prescreening survey to determine eligibility. All participants were regular cannabis users with no evidence of life problems stemming from their drug use (based on a modified version of the Drug Abuse Screening Test [26]), were not in treatment or ever recommended for treatment, and had no signs of psychosis (based on a brief 7-item screener [27]). We included heavy users to ensure the sample was representative of those most likely to drive while under the influence.

#### Protocol

Participants were run in groups of up four at a time, with two groups run per day. To ensure no one drove home after using cannabis, taxi rides home were provided for individuals who could not arrange to be picked up by friends or family after the experimental session. Participants were asked to legally purchase cannabis (in flower form) to bring to the study session, along with a dosing implement (e.g., pipe, bong). The use of commercially available cannabis strains with potencies that the public actually uses—rather than those strains provided by the National Institute for Drug Abuse—helped ensure greater representativeness and generalizability of findings.

Participants completed four blocks of attentional and psychomotor tests: a baseline block, which was followed by a cannabis dosing session (during which participants took up to five hits of the cannabis they purchased legally for the research), and then three additional post-dosing blocks (each starting approximately 1 hour apart). Data collection from the same participants over time natural created variability in cannabinoid concentrations, which was used to predict task performance. We asked participants to refrain from cannabis use 5 hours prior to data collection to ensure that all participants had low and high measured concentrations, but unlike a traditional experimental dosing studies, our design did not require complete sobriety at baseline (analysis focused on THC and CBD concentrations, not performance at different times).

A saliva sample was collected from each participant (using Quantisal™ collection tubes) at the start of each block (baseline and the three follow-up testing blocks) and assayed for concentrations of THC and CBD by Immunalysis Corporation) using the same menhod previously described in Study 1.

#### Measures

Each block of testing included (in randomized order) the Sustained Attention to Response Task (SART) of vigilance and impulse control [36], the Tower of London (ToL) task of executive function and planning [37], and the Pursuit Rotor task of visual-tracking skills and coordination [38]. Tests were administered using Inquisit™ software. The key dependent measures consisted of performance on the SART, the ToL test, and the Pursuit Rotor task.

The **SART** involved displaying a rapid, pseudo-random sequence of numbers 1 to 9; participants were instructed to respond as quickly as possible by pressing the space bar when a number appeared. An exception was that when the number 3 appeared participants were instructed to withhold their response. The dependent measure was participants’ mean response latency to pressing the space bar during the series of trials immediately before successfully withholding a response to a 3—how fast can participants respond to the correct cues and still avoid responding to a 3. Participants with diminished ability to sustain focus and/or control impulses need to respond more slowly—i.e., they have *larger* response latencies on space bar responses in order to effectively withhold that response at the right time.

The **ToL** task involved a computerized graphic of three poles with colored “donuts” placed on them. Participants were instructed to rearrange the donuts to match a target arrangement (shown in an image) within a set number of moves. The task requires planning ahead and executive functioning in order to efficiently match the target. Scores were assigned over 12 trials such that successfully matching the standard in the fewest moves generated more points. Participants with diminished executive functioning should have lower ToL scores.

The **Pursuit Rotor** task involved a computerized dot moving in a circle at a fixed speed on the screen (imagine a dime spinning around on the edge of a turntable). Participants were instructed to hover the mouse pointer on top of dot as it moved. The dependent measure was the percentage of time hovering on top of the image. Participants with diminished tracking and coordination would have a lower percentage.

### Study 2: Results

#### Sample

Twenty-four regular cannabis users were recruited out of 55 who indicated interest in the research. Of the participants, slightly more than half (n = 13) were male with ages ranging from 21 to 56 years (median 32). The majority (n = 16) of participants were White and non-Hispanic; four were African American, three were Hispanic (White), and one was Native American. Participants included heavy cannabis users. The majority (n = 17) used multiple times per day, five used daily or almost daily, and two used multiple times per week. Participants were run in groups of up to four at a time, and two groups were run per day. THC and CBD concentrations assayed from participants’ oral fluid samples for each block are provided in Table 7. Approximately half of the cannabis used by participants contained no measurable CBD.

**Table 7.** Descriptive Information of Cannabinoid Concentrations Across Blocks for Pilot Study.

|  | **Baseline Block** | **Post-Dosing Block 1** | **Post-Dosing Block 2** | **Post-Dosing Block 3** |
| --- | --- | --- | --- | --- |
| **THC** | *N* = 24<br>Mean = 47.7 (88.7)<br>Minimum = 0.0<br>Maximum = 417 | *N* = 24<br>Mean = 554.0 (550.3)<br>Minimum = 22<br>Maximum = 1734 | *N* = 24<br>Mean = 214.7 (347.2)<br>Minimum = 6<br>Maximum = 1467 | *N* = 24<br>Mean = 103.7.7 (172.0)<br>Minimum = 4<br>Maximum = 840 |
| **CBD** | *N* = 24<br>Mean = 0.8 (2.3)<br>Minimum = 0<br>Maximum = 9 | *N* = 24<br>Mean = 14.2 (27.4)<br>Minimum = 0<br>Maximum = 103 | *N* = 24<br>Mean = 7.2 (17.0)<br>Minimum = 0<br>Maximum = 72 | *N* = 24<br>Mean = 2.2 (4.8)<br>Minimum = 0<br>Maximum = 17 |

#### Analytic approach

Analyses were conducted using generalized linear mixed modeling in SAS 9.2. Like the previously described driving study, we predicted performance on the computerized tasks from THC and CBD scores obtained from participants’ oral fluid samples, looking at the same subjects at different drug levels at different times during a laboratory session. Unlike the driving study, we used quantitative concentration of CBD (ng/ml) rather that a dichotomous presence or absence indicator. We did not log transform THC or CBD concentrations obtained in the laboratory, as the controlled measurement allowed for better data distributions. Test block was included as a control variable to help account for generic time effects. Self-reported frequency of cannabis use collected during prescreening was used as a covariate (the item was measured on a 7-point scale from “a few times per year” to “several times per day”). Since each participant provided multiple oral fluid and performance test scores over time, we modeled “subject” as a random effect to accommodate within-subject correlations.

#### Tests of hypotheses

##### Sustained Attention to Response Task

The SART produced data on response latencies (mean = 392.4 ms, SD = 131.3)—how quickly participants responded to stimuli and still avoided making errors. Larger response latencies indicated greater impairment (i.e., impaired participants needed to go more slowly to avoid errors). Analysis revealed a statistically significant THC x CBD interaction and a main effect of CBD concentrations on response latencies (see Table 8).

**Table 8.**
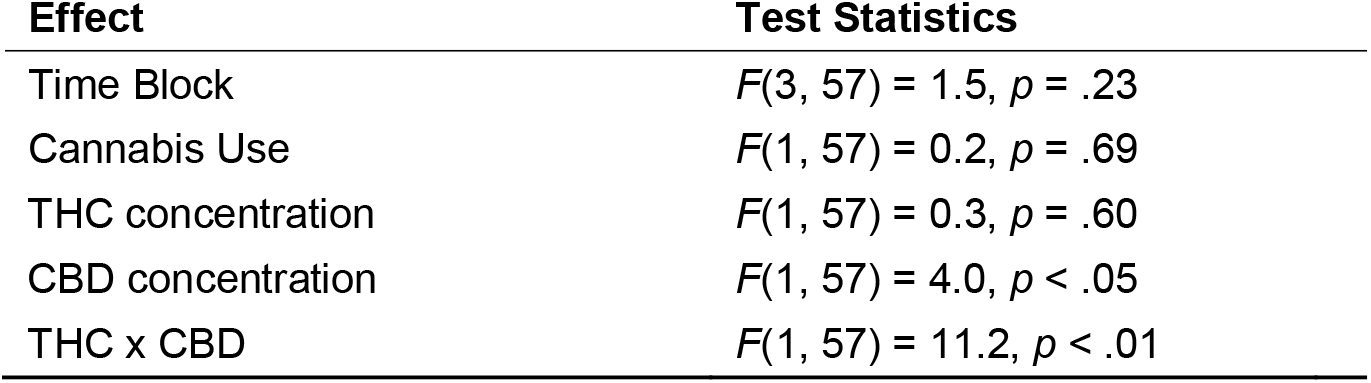
Analysis of the SART Results from THC x CBD.

| Effect | Test Statistics |
| --- | --- |
| Time Block | $F(3, 57) = 1.5, p = .23$ |
| Cannabis Use | $F(1, 57) = 0.2, p = .69$ |
| THC concentration | $F(1, 57) = 0.3, p = .60$ |
| CBD concentration | $F(1, 57) = 4.0, p < .05$ |
| THC x CBD | $F(1, 57) = 11.2, p < .01$ |

The interaction between continuous variables was interpreted using the method outlined by Aiken and West [39] for probing interactions in regression. In Fig 6, we illustrate the linear-by-linear interaction effect, displaying slopes for THC concentrations against reaction time scores on the SART, with separate slopes for low and high CBD concentrations. For this study, per the method described by Aiken and West [39], we defined low CBD as zero and set the high CBD to a value of 50, approximately one standard deviation above the block 1 mean. The pattern displayed in Fig 6 was created by solving the regression equation (using the ESTIMATE command in SAS) between THC and reaction time scores when CBD is 0 and when CBD is 50 to reflect low and high CBD levels. We could have computed slopes at any CBD values, but slopes at other CBD values can be interpolated from the figure. The goal of the research was not to identify specific THC or CBD thresholds but rather to confirm that the skill impairment varied not only as a function of THC, but as a function of CBD as well.

**Figure 6.**
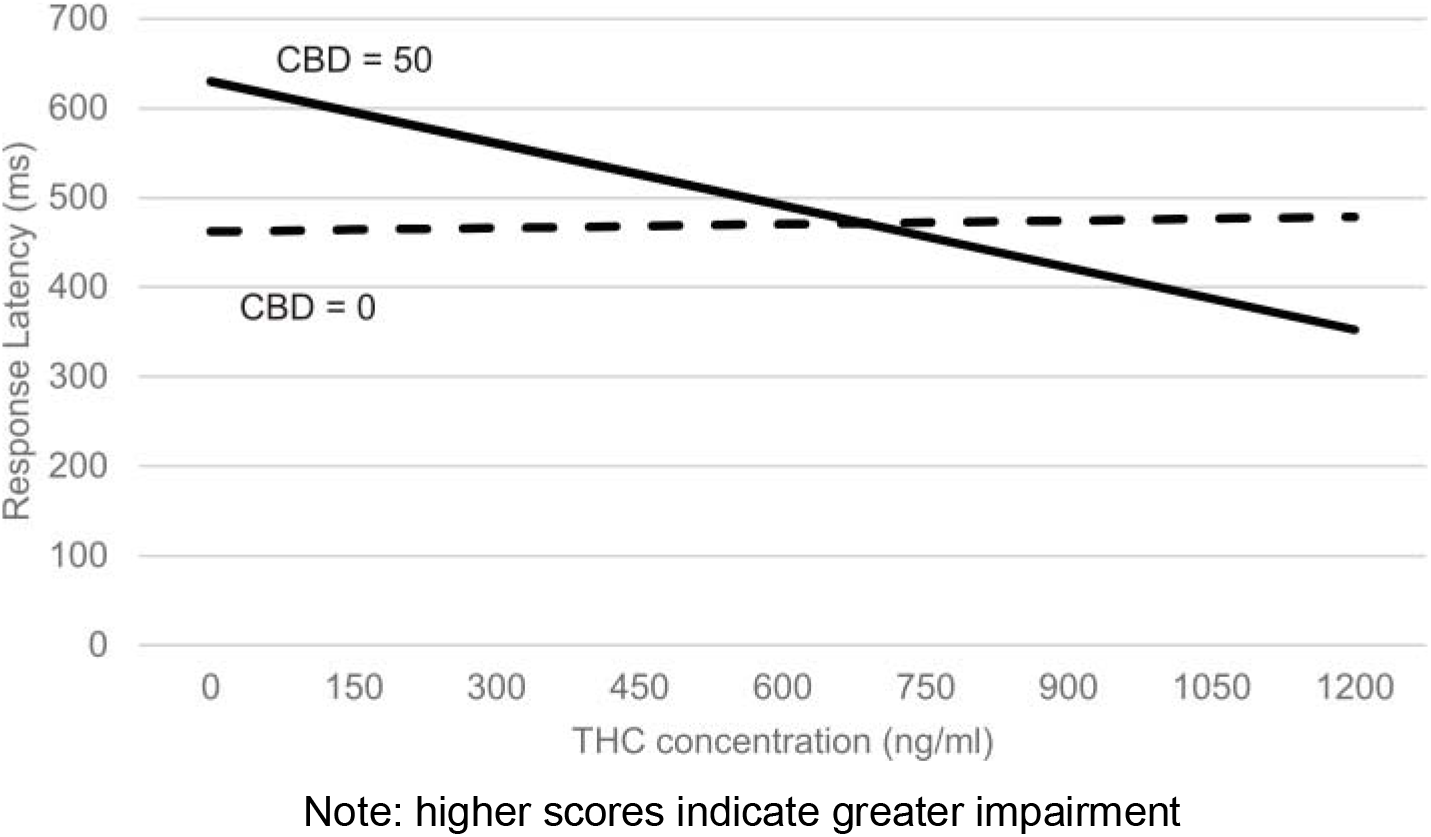
Sustained Attention to Reaction Task (SART) as a Function of THC and CBD

Accordingly, reaction times were slower (larger) when participants had relatively high levels of CBD in their systems, however, only when THC concentrations were also relatively low or moderate. When CBD was set to 0 ng/ml, there was little relationship between THC and task performance. When CBD was set to 50 ng/ml, the relationship between THC and impairment was negative. This pattern matches the pattern that was observed in the driving study regarding the standard deviation of X-axis acceleration (Fig 4) and vehicle speed (Fig 5).

##### Tower of London task

ToL scores (mean = 29.4, SD = 4.9) reflect executive functioning, with higher scores indicating better planning—solving the set of problems in fewer moves [37]. However, analysis of ToL scores revealed only a marginally significant THC x CBD interaction (p = .08), but no significant main effects after removing the interaction effect. These effects were not interpreted further. There were significant main effects for time block and cannabis use (both p-values < .0). Overall, performance scores improved over time (block), as participants learned to solve the different examples. Scores also were better to the extent that participants were more frequent cannabis users (B = -2.4).

The Pursuit Rotor task outcome (mean = 45.4%, SD = 19.8) reflected the amount of time participants were on target during the tracking task. Analysis revealed no significant main effects of interactions on Pursuit Rotor task scores.

### Study 2: Discussion

On the SART task, participants with high levels of CBD but low to moderate levels of THC exhibited worse performance. This study complements and confirms some findings from the driving study (Figs 4 and 5), demonstrating that the combination of lower THC but higher CBD predicted driving. Given that impairment was observed only when THC concentrations were low suggests that CBD operates through a wholly different path of impairment than THC, for example via mental sedation versus through psychoactive effects. The study produced no evidence of impairment due to high THC concentrations. However, given that our participants were mostly very heavy cannabis users, this is not terribly surprising. Previous research has found that heavy users sometimes are tolerant to the impairing effects of THC [34, 40].

Like the driving study, this research was epidemiological, not experimental; there was no control over assignment of high versus low CBD cannabis. Participants self-selected their cannabis strains and thus individual personality characteristics invariably could be confounded with CBD concentrations.

## Study 3

A third study was conducted to replicate and expand upon Study 2 and further confirm that evidence generated from the driving study. Like Study 2, the sample consisted primary of heavy cannabis users. While we still were unable to legally acquire, control and dose participants with specific strains, Study 3 included additional control features and more measures to add to the rigor of the design. We do not consider this to be controlled dosing study where cannabis consumption is experimentally manipulated, but rather a version of laboratory epidemiology.

### Study 3: Materials and Methods

The Institutional Review Board of the Pacific Institute for Research and Evaluation approved all aspects of the Study 3 research described herein, including the protocol for participants self-dosing with cannabis.

#### Protocol

The research took place in Denver, Colorado. Participants for the main study were primarily heavy cannabis users recruited using an approach similar to the one described in Study 2. In this study, rather than allowing participants to bring in their preferred cannabis, subjects were asked (randomly determined) to purchase and bring to the study one of three broad categories of commercially available product. Participants also each took part in two experimental sessions using a different type of cannabis each time. During each experimental session, participants completed four blocks of computerized attentional and psychomotor tests: an unrecorded practice block (to help reach stable performance levels), a recorded baseline block followed by the cannabis dosing session, and then two post-dosing blocks. An oral fluid sample was collected during the baseline and two post-dosing blocks. Unlike the pilot study participants, who were limited to only five hits of cannabis, participants in the main study could smoke up to 1 gram of their assigned cannabis. Research assistants weighed participants’ cannabis before and after dosing to determine the amount consumed.

For a given experimental session, participants were randomly assigned to either a high THC/low CBD strain (12-18% THC, <1% CBD), a balanced 1:1 strain (5-10% THC, 5-10% CBD), or a CBD-dominate stain (e.g., ~4% THC, 10% CBD). The purpose of this assignment was not to allow causal statements—which was not possible because we could not actually control or administer the cannabis—but to help ensure sufficient variability in THC and CBD concentrations. Across both sessions participants were asked to purchase two of the three broad types of cannabis. Once again, the research focused on measured cannabinoid concentrations rather than a direct comparison of specific strain potencies, as we did not have sufficient control over the cannabis to ensure high fidelity of assignment.

Each testing block included five cognitive and psychomotor tests provided by *Cambridge Cognition™* and administered to participants on touch-sensitive tablets (an external press-pad was used for some of the tests). The tests tapped into skills ostensibly related to safe driving and included: (a) Rapid Visual Processing—a test of sustained attention; (b) Reaction Time—a measure of psychomotor speed; (c) Motor Screening—a general sensorimotor assay of hand-eye coordination, response speed, and accuracy; (d) Attention Switching—a test of focus, attention, and top-down cognitive control processes; and (e) Information Sampling—a test of impulsivity and decision making. The test order was randomized among subjects but consistent within-subjects to ensure consistent time from the oral fluid collection.

Oral fluid samples (collected during baseline and the two follow-up periods) were assayed for quantitative concentrations of cannabinoids using the same approach as described in the pilot study. Self-reported frequency of both cannabis use was collected during the prescreening survey and used in analysis as covariates.

### Study 3: Results

#### Sample

A total of 91 individuals completed the online prescreening survey; 40 were deemed eligible and agreed to participate. However, only 35 (21 men and 14 women) completed both sessions of the experiment. Ages ranged from 22 to 61 years with a median age of 41. The majority (n = 20) were White and non-Hispanic; seven were African American, six were Hispanic (White), and two were Native American. Most (n = 20) used cannabis multiple times per day, nine used daily or almost daily, and six used multiple times per week. Participants tended to be infrequent drinkers, consuming alcohol an average of 2.5 days (SD = 3.9) during the past 30. Mean THC and CBD levels, across blocks, are provided in Table 9. Note that THC and CBD levels are much higher in this study than in Study 1 because the strains used were relatively more potent and because participants were free to consume up to 1 gram of cannabis.

**Table 9.** Descriptive Information of Cannabinoid Concentrations Across Blocks for Study 3.

|  | **Baseline Block** | **Post-Dosing Block 1** | **Post-Dosing Block 2** |
| --- | --- | --- | --- |
| **THC** | $N = 36$<br>Mean = 116.2 ($SD = 321.2$)<br>Minimum = 0.0<br>Maximum = 417 | $N = 36$<br>Mean = 793.8 ($SD = 852.6$)<br>Minimum = 17<br>Maximum = 2949 | $N = 36$<br>Mean = 152.9 ($SD = 216.0$)<br>Minimum = 3<br>Maximum = 1232 |
| **CBD** | $N = 36$<br>Mean = 0.7 ($SD = 2.2$)<br>Minimum = 0<br>Maximum = 13 | $N = 36$<br>Mean = 136.5 ($SD = 322.9$)<br>Minimum = 0<br>Maximum = 1683 | $N = 36$<br>Mean = 15.0 ($SD = 33.5$)<br>Minimum = 0<br>Maximum = 202 |

Four cases with outlier THC scores were removed.

#### Data reduction

The five computerized tests produced 18 distinct outcomes measures. Instead of analyzing each outcome separately and incurring the risk of making Type I errors, we subjected the collection of outcomes to a principle components analysis (PCA, with direct oblim rotation) and generated reliable factor scores that reflected skills or sets of skills common among the measures. Our analysis focused only on the first extracted factor—which accounted for 20.4% of the variance among the items—as the dependent measure. This factor was defined largely by delayed response latencies on the Attention Switching task (loading = .808) and Reaction Time test (.685), as well as slow movement time on the Reaction Time test (.620). Congruency and switching costs from the Attention Switching task also loaded positively (.563 and .592, respectively). Higher factor scores definitively indicated greater impairment, as they reflect the costs (slower responses) due to being able to focus in light of distractions and to shift focus when needed.

#### Cannabis consumption

The difference between pre-smoking and post-smoking cannabis weights was examined. On average, participants consumed two thirds of a gram (mean = .667, SD = .51) of cannabis. Although there certainly was variability in consumption among participants, the weight of cannabis consumed did not differ significantly as a function of the three strain conditions (*p* = .85). Further, paired sample t-test found no significant differences in amount of cannabis consumed by the same participants between the two sessions (*p* = .22).

#### Analytic approach

As with Study 2, the attention-impaired factor score was analyzed using generalized linear mixed modeling. The main effects of THC and CBD concentrations, as well as the THC x CBD interaction, were the primary predictors. Block (baseline, post-dosing 1, and post-dosing 2), session, and the Block x Session interaction were included as control variables. “Subject” was modeled as a random effect to accommodate non-independent responses given that each participant took part in multiple trials. Frequency of cannabis use and frequency of alcohol consumption were modeled as covariates.

#### Tests of hypotheses

We eliminated from analysis cases associated with four outlier high THC scores (>3000 ng/ml), which were discontinuous within the distribution. Including these cases, however, had minimal bearing on the results.

Results of the generalized linear model analysis are shown in Table 10. There was a significant effect of time block, with improving performance over time (controlling for drug concentrations) but no main effect of session and no Block x Session interaction. Self-reported frequency of cannabis use was significantly but positively related to the reaction time factor (B = 0.44); subjects who used cannabis more frequently demonstrated overall worse (slower) performance (again, controlling for other factors in the model). The main effect of THC was statistically significant (B = -0.0002), suggesting improved overall performance with increasing THC concentrations. Given that both studies #1 and #2 produced statistically significant THC x CBD interactions, we interpreted the linear by linear THC x CBD interaction (p = .07) as significant given that a one-tailed test is justified.

**Table 10.** Analysis of the Test Results from THC and CBD.

| **Effect** | **Test Statistics** |
| --- | --- |
| Time Block | $F(2, 118) = 3.9, p < .05$ |
| Session | $F(1, 63) = 1.4, p = .25$ |
| Time Block x Session | $F(2, 118) = 1.6, p = .20$ |
| Cannabis Use | $F(1, 118) = 6.7, p < .05$ |
| THC concentration | $F(1, 118) = 5.3, p < .05$ |
| CBD concentration | $F(1, 118) = 2.1, p = .16$ |
| THC x CBD | $F(1, 118) = 3.4, p = .07$ |

The interaction was probed using the method outlined by Aiken and West [39] and is depicted in Fig 7. We solved regression equations that predicted task performance from THC at CBD values of 0 and 500 (approximately 1 standard deviation above the block 1 mean) (see Table 9). THC and CBD concentration scores were notably higher than in the pilot study due to use of more potent strains and/or greater amounts of cannabis consumed.

**Figure 7.**
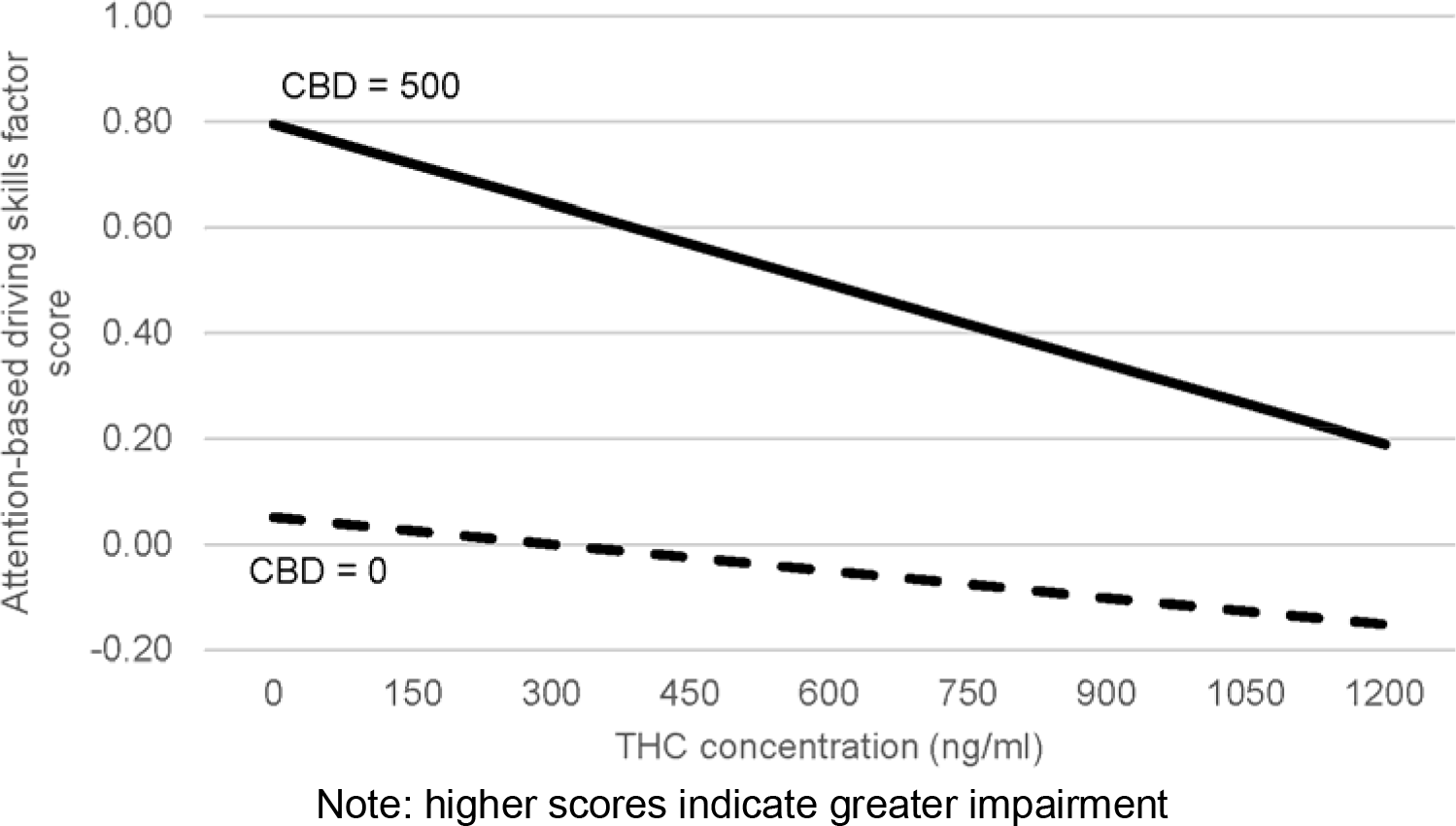
Attention-Based Driving Skills a Function of THC and CBD

As shown in Fig 7, higher CBD concentrations were associated with more impaired performance (i.e., increased reaction times)—with exacerbated effects when THC concentrations also were lower. At high THC concentrations, however, CBD concentrations were relatively unrelated to task performance. THC concentrations were unrelated to task performance at low CBD levels, and at higher CBD levels, increased THC was associated with increased task performance (reduced reaction times).

### Study 3: Discussion

The results on the composite skill score mirror those findings from Study 2 and observed in Study 1; impaired performance was observed when CBD concentrations where high and THC concentrations were low. The study protocol—assigning participants to cannabis strain categories and relying on doubly within subject comparisons (over time and within sessions)— was designed to minimize alternative explanations. Nevertheless, we were careful not to frame the results as causal effects.

## General discussion

In three studies, driving performance or performance on driving-related skills was predicted by cannabinoid concentrations, with CBD the apparent driver of effects. In every case, riskier driving was associated with higher CBD levels. Greater prevalence of risky driving (i.e., elevated g-force events, which have been linked to actual crash risk) was observed among drivers who tested higher in THC but also were CBD positive. Greater variability in speed (i.e., lack of speed control), greater vehicle speed, and impairment on the attention-based psychomotor tasks (Studies 2 and 3) were observed when CBD was higher but THC concentrations were low. Greater variability in lateral vehicle control was associated with being CBD positive (versus CBD negative), regardless of THC. In general, high THC levels in the absence of CBD was not associated with impaired performance. This should not be particularly surprising, however, given that our sample consisted of very heavy cannabis users who may have developed drug tolerance and demonstrated less impairment due to THC [34, 40].

This research was not experimental; participants used commercially available cannabis strains, and we could not legally manage the product prior to dosing nor administer the dose to participants in a highly controlled fashion. The driving study (Study 1) observed naturalistic cannabis use over time, using a within-subjects design to capture (potentially) different strains and concentrations within the same drivers over time. Further, Study 3 assigned subjects to broad strain types in an effort to divorce personality from drug selection. Nevertheless, it is possible that individuals with specific driving styles, risk-taking tendencies, etc., would naturally choose one type of cannabis over other (e.g., sativa versus indica). Additionally, with the exception of Study 3, the cannabis consumed (based on measured drug concentrations) was predominately CBD-free. A better balance of low versus high CBD strains would have been preferred.

The research did not include any placebo control. A placebo control was not applicable in the driving study and not legally possible in the two laboratory studies; however, it also is not typically used for epidemiological research. Placebo control conditions often are used to rule out learning or fatigue effects; however, our key findings cannot easily be explained by those because our primary conclusions concern impairment as a combined function of THC and CBD. For the results of the laboratory studies (see Figs 6 and 7) to be attributed to fatigue, for example, one would need to explain why only subjects with high CBD and low THC experienced fatigue. The fatigue effect explanation is not relevant to the driving study.

Finally, even in a true, experimental controlled dosing paradigm it might be difficult to make causal statements regarding compounds within a drug as complex of cannabis. With hundreds of cannabinoids and terpenes within cannabis, it is possible that random assignment across strain in fact varies along more than one compound. For example, while several experimental studies have linked high-CBD cannabis to the experience of lethargy and sedation [18, 19], others have argued that pharmaceutical CBD, in fact, is not sedating [23]. Rather, they argue high-CBD cannabis contains the terpene *myrcene*, which is a defining feature of *cannabis indica* and the sedated “body high” typical of high CBD strains. In other words, although we associated high CBD concentrations with driving impairment, it is possible, at least, that CBD i¡ only a proxy of myrcene, and that compound is the causal agent. To our knowledge, there currently is no opportunity to assay myrcene concentrations from oral fluid.

In summary, our research found that cannabis with CBD produced different driving patterns, and different degrees of driving skill impairment, than cannabis lacking CBD. Risky driving (elevated g-force events) was associated with high THC concentrations and positive CBD results. Other indices of driving and driving-skill impairment were associated with higher CBD and low THC concentrations, or simply with higher CBD levels. More research is needed to explore the role of compounds other than THC on driving impairment, including true experimental studies as well as epidemiological relative risk research. The results described herein—evidence that THC alone cannot explain driving impairment—provides justification for future research efforts in this area.

## References

1. Berning A, Compton R, Wochinger K. Results of the 2013-2014 National Roadside Survey of Alcohol and Drug Use by Drivers. Washington, DC:National Highway Traffic and Safety Administration; 2015.

2. Johnson MB, Kelley-Baker T, Voas RB, Lacey JH. The prevalence of cannabis-involved driving in California. Drug Alcohol Depend. 2012;12(1-3):105–9. doi:http://dx.doi.org/10.1016/j.drugalcdep.2011.10.023. PubMed Central PMCID: PMCPMCID: PMC3755617.

3. Masten SV, Guenzburger GV. Changes in driver cannabinoid prevalence in 12 US states after implementing medical marijuana laws. J Safety Res. 2014;50:35–52.

4. Salomonsen-Sautel S, Min S-J, Sakai JT, Thurstone C, Hopfer C. Trends in fatal motor vehicle crashes before and after marijuana commercialization in Colorado. Drug Alcohol Depend. 2014;140:137–44. doi:10.1016/j.drugalcdep.2014.04.008. PubMed Central PMCID: PMCPMCID: PMC4068732.

5. Hartman R, Huestis M. Cannabis effects on driving skills. Clin Chem. 2013;59(3):478–92. PubMed Central PMCID: PMCPMCID:PMC3836260.

6. Capler R, Bilsker D, Van Pelt K, MacPherson D. Cannabis Use and Driving: Evidence Review. Vancouver, BC: Canadian Drug Policy Coalition (CDPC), Simon Fraser University, 2017.

7. Hels T, Bernhoft IM, Lyckegaard A, Houwing S, Hagenzieker M, Legrand S, et al. Risk of injury by driving with alcohol and other drugs. 2011 [cited August 2012]. In: DRUID (Driving Under the Influence of Drugs, Alcohol and Medicines) Deliverable D235 [Internet]. Bergisch-Gladbach, Germany: Federal Highway Research Institute, [cited August 2012]. Available from: http://www.druid-project.eu/Druid/EN/deliverales-list/downloads/Deliverable_2_3_5.pdf?_blob=publicationFile.

8. Lacey JH, Kelley-Baker T, Berning A, Romano E, Ramirez A, Yao J, et al. Drug and alcohol crash risk: A case-control study. Washington, DC: National Highway Traffic Safety Administration; 2016.

9. Gjerde H, Strand MC, Mørland J. Driving under the influence of non-alcohol drugs—An update. Part I: Epidemiological studies. Forensic Sci Rev. 2015;27(2):89–113.

10. Aizpurua-Olaizola O, Soydaner U, O_ztu_rk E, Schibano D, Simsir Y, Navarro P, et al. Evolution of the cannabinoid and terpene content during the growth of cannabis sativa plants from different chemotypes. J Nat Prod. 2016;79(2):324–31.

11. Joy J, Watson S, Benson J, editors. Marijuana and medicine: Assessing the science base. Washington, DC: National Academy of Sciences Press; 1999.

12. Pearce DD, Mitsouras K, Irizarry KJ. Discriminating the effects of Cannabis sativa and Cannabis indica: A web survey of medical cannabis users. Journal of Alternative and Complementary Medicine. 2014;20(10):787–91. Epub 2014/09/06. doi:10.1089/acm.2013.0190. PubMed PMID:25191852.

13. Hazekamp A, Fischedick JT. Cannabis-from cultivar to chemovar. Drug Testing and Analysis. 2012;4(7-8):660–7. doi:doi:10.1002/dta.407.

14. Hillig KW, Mahlberg PG. A chemotaxonomic analysis of cannabinoid variation in Cannabis (Cannabaceae). American Journal of Botany. 2004;91(6):966–75.

15. Moore TH, Zammit S, Lingford-Hughes A, Barnes TR, Jones PB, Burke M, et al. Cannabis use and risk of psychotic or affective mental health outcomes: A systematic review. Lancet. 2007;370(9584):319–28.

16. Zuardi AW. History of cannabis as a medicine: A review. Rev Bras Psiquiatr. 2006;28(2):153–7.

17. Morgan CJ, Curran HV. Effects of cannabidiol on schizophrenia-like symptoms in people who use cannabis. Br J Psychiatry. 2008;192(4):306–7. doi:doi:10.1192/bjp.bp.107.046649.

18. de Souza Crippa JA, Zuardi AW, Garrido GE, Wichert-Ana L, Guarnieri R, Ferrari L, et al. Effects of cannabidiol (CBD) on regional cerebral blood flow. Neuropsychopharmacol. 2004;29(2):417–26.

19. Zhornitsky S, Potvin S. Cannabidiol in humans—The quest for therapeutic targets. Pharmaceuticals. 2012;5(5):529–52.

20. Martin A. Major compounds in cannabis and the impact to your high: Weedist; 2012. Available from:http://www.weedist.com/2012/06/major-compounds-in-cannabis-and-the-impact-to-your-high.

21. Frank M, Rosenthal E. The Marijuana Grower’s Guide. London: Atlantic Books; 1988.

22. Russo EB. Taming THC: Potential cannabis synergy and phytocannabinoid-terpenoid entourage effects. British Journal of Pharmacology. 2011;163(7):1344–64. doi:10.1111/j.1476-5381.2011.01238.x. PubMed PMID: PMC3165946.

23. Piomelli D, Russo EB. The Cannabis sativa versus Cannabis indica debate: An interview with Ethan Russo, MD. Cannabis Cannabinoid Res. 2016;1(1):44–6.

24. Evans L. Traffic Safety: Science Serving Society; 2004.

25. Klauer SG, Dingus TA, Neale VL, Sudweeks JD, Ramsey DJ. The Impact of Driver Inattention on Near-Crash/Crash Risk: An Analysis Using the 100-Car Naturalistic Driving Study Data. Washington, DC: National Highway Traffic Safety Administration, 2006.

26. Gavin DR, Ross HE, Skinner HA. Diagnostic validity of the DAST in the assessment of DSM-III drug disorders. Br J Addict. 1989;84(3):301–7.

27. Degenhardt L, Hall W, Korten A, Jablensky A. Use of a brief screening instrument for psychosis: Results of an ROTC analysis. New South Wales, Sydney, Australia: National Drug and Alcohol Research Centre, 2005 NDARC Technical Report No. 210.

28. Lacey JH, Kelley-Baker T, Furr-Holden D, Voas RB, Moore C, Brainard K, et al. 2007 National Roadside Survey of Alcohol and Drug Use by Drivers. Washington, DC: U.S. Department of Transportation, National Highway Traffic Safety Administration; 2009.

29. Kelley-Baker T, Lacey JH, Berning A, Ramirez A, Moore C, Brainard K, et al. 2013-2014 National Roadside Study of alcohol and drug use by drivers: Methodology. Washington, DC: National Highway Traffic Safety Administration; 2016.

30. Simons-Morton BG, Zhang Z, Jackson JC, Albert PS. Do elevated gravitational-force events while driving predict crashes and near crashes? American Journal of Epidemiology. 2012;175(10):1075–9. doi:10.1093/aje/kwr440.

31. Bagdadi O, Varhelyi A. Jerky driving‐‐An indicator of accident proneness? Accid Anal Prev. 2011;43(4):1359–63. Epub 2011/05/07. doi:10.1016/j.aap.2011.02.009. PubMed PMID:21545866.

32. Sewell RA, Poling J, Sofuoglu M. The effect of cannabis compared with alcohol on driving. Am J Addict. 2009;18(3):185–93. Epub 2009/04/03. doi:10.1080/10550490902786934. PubMed PMID: 19340636; PubMed Central PMCID: PMCPMCID:PMC2722956.

33. Schwope DM, Bosker WM, Ranaekers JG, Gorelick DA, Huestis MA. Psychomotor performance, subjective and physiological effects and whole blood Δ9-tetrahydrocannabinol concentrations in heavy, chronic cannabis smokers following acute smoked cannabis. J Anal Toxicol. 2012;36(6):405–12.

34. Ramaekers JG, Theunissen EL, De Brouwer M, Toennes SW, Moeller MR, Kauert G. Tolerance and cross-tolerance to neurocognitive effects of THC and alcohol in heavy cannabis users. Psychopharmacol. 2011;214(2):391–401.

35. Hartman RL, Brown TL, Milavetz G, Spurgin A, Pierce RS, Gorelick DA, et al. Cannabis effects on driving lateral control with and without alcohol. Drug Alcohol Depend. 2015;154:25–37.

36. Robertson IH, Manly T, Andrade J, Baddeley BT, Yiend J. 'Oops!': performance correlates of everyday attentional failures in traumatic brain injured and normal subjects. Neuropsychologia. 1997;35(6):747–58.

37. Krikorian R, Bartok J, Gay N. Tower of London procedure: a standard method and developmental data. Journal of clinical and Experimental Neuropsychology. 1994;16(6):840–50.

38. Adams JA. Warm-up decrement in performance on the pursuit-rotor. The American Journal of Psychology. 1952;65(3):404–14.

39. Aiken LS, West SG. Multiple regression: Testing and interpreting interactions. Thousand Oaks, CA: Sage; 1991.

40. Schwope DM, Bosker WM, Ranaekers JG, Gorelick DA, Huestis MA. Psychomotor performance, subjective and physiological effects and whole blood Δ9-tetrahydrocannabinol concentrations in heavy, chronic cannabis smokers following acute smoked cannabis. Journal of Analytic Toxicology. 2012;36(6):405–12.

